# Amyloid precursor protein induces reactive astrogliosis

**DOI:** 10.1101/2023.12.18.571817

**Authors:** Gretsen Velezmoro Jauregui, Dragana Vukić, Isaac G. Onyango, Carlos Arias, Jan S. Novotný, Kateřina Texlová, Shanshan Wang, Kristina Locker Kovačovicova, Natalie Polakova, Jana Zelinkova, Maria Čarna, Valentina Lacovich Strašil, Brian P. Head, Daniel Havas, Martin Mistrik, Robert Zorec, Alexei Verkhratsky, Liam Keegan, Mary O’Connel, Robert Rissman, Gorazd B. Stokin

## Abstract

We present *in vitro* and *in vivo* evidence demonstrating that Amyloid Precursor Protein (APP) acts as an essential instigator of reactive astrogliosis. Cell-specific overexpression of APP in cultured astrocytes led to remodelling of the intermediate filament network, enhancement of cytokine production and activation of cellular programs centred around the interferon (IFN) pathway, all signs of reactive astrogliosis. Conversely, APP deletion in cultured astrocytes abrogated remodelling of the intermediate filament network and blunted expression of IFN stimulated gene (ISG) products in response to lipopolysaccharide (LPS). Following traumatic brain injury (TBI), mouse reactive astrocytes also exhibited an association between APP and IFN, while APP deletion curbed the increase in glial fibrillary acidic protein (GFAP) observed canonically in astrocytes in response to TBI. Thus, APP represents a molecular inducer and regulator of reactive astrogliosis.

## INTRODUCTION

Astrocytes are homeostatic cells of the central nervous system (CNS) that provide support and regulate various aspects of the functional activity of the nervous tissue being fundamental elements of the brain active milieu^1^. In particular, astrocytes contribute to the regulation of synaptic transmission ^2–5^, regulation of energy metabolism ^6–9^ and maintenance of the blood brain barrier ^10–13^. Brain lesions trigger reactive astrogliosis which, together with microgliosis and mobilisation of oligodendrocyte precursor cells, represents evolutionary conserved defensive response aimed at protection and postlesional restoration of the nervous tissue from various forms of pathological stressors and insults ^14,15^ Reactive astrogliosis proceeds through complex, still poorly understood, reorganisation of astrocytic biochemistry, morphology and physiology to create context- and disease-specific reactive phenotypes ^16,17^. Reactive astrogliosis is linked to the activation of several cellular programs and second messenger signalling pathways ^15,18^ that still remain to be fully elucidated ^19–23^. In particular, reactive astrocytes up-regulate synthesis and release of pro-inflammatory proteins such as cytokines ^24,25^ and antigen-presenting molecules ^26^. Molecular pathways which trigger and regulate the progression of reactive astrogliosis remain however largely unknown. Accumulating evidence indicates that various insults instigating reactive astrogliosis increase expression of the amyloid precursor protein (APP), an astrocyte resident type I integral membrane protein involved in many physiological and pathophysiological processes ^27–31^. For example, intraperitoneal injection of lipopolysaccharide (LPS), an archetypal pathogen-associated molecular pattern, increases APP levels in astrocytes in mice and rats ^32–34^. Similar increase was observed after intraventricular injection of LPS ^35^ or by adding LPS to astrocyte culture media ^36^. Treatment with cytokines such as interleukin 1 (IL1) ^36–40^, tumour necrosis factor β (TNFβ) ^38,39,41^, tumour necrosis factor α (TNFα) ^36^ and interferon γ (IFNγ) ^36,37^ increased APP levels in astrocytes both *in vitro* and *in vivo*. Astrocytic APP levels were increased in many pathological conditions including ischaemia ^42^, brain trauma ^43,44^ heat shock^45^, cuprizone induced demyelination ^46^, quinolinic acid excitotoxicity ^47^ and hyperammonaemia^48,49^ as well as following increase in cytoplasmic cyclic adenosine monophosphate (cAMP) ^50^. In all of these conditions, astrocytes frequently generate molecules involved in inflammation like cytokines. ^51–53^ There is little knowledge, however, how molecular mechanisms exactly orchestrate the acquisition of this reactive state of astrocytes.

The evidence that all these stressors and conditions do not only induce reactive astrogliosis, but invariably increase expression levels of APP in astrocytes led us to hypothesise that APP plays a pivotal role in the transition of astrocytes into a reactive state. To test this hypothesis, we utilized molecular, biochemical and immunohistochemical approaches in cell culture and in a mouse model of TBI to rigorously investigate the role of 770 amino acid long APP ^31,38^ in the initiation of reactive astrogliosis. We found that overexpression of this APP isoform in astrocytes promotes, while deletion of APP abrogates reactive astrogliosis, thus confirming that APP represents a molecular switch initiating reactive astrogliosis. Astrocytic APP may thus be a valuable target in the regulation of the response of astroglia to pathological conditions.

## RESULTS

### APP triggers reactive astrocyte morphology

As alluded before, previous studies showed that stressors inducing reactive astrogliosis at the same time also increase APP expression ^34–36^. This raised the question of whether increased APP levels induce reactive state of the astrocytes in the absence of stressors. As the first step, we reproduced previously reported effects of LPS on the GFAP-immunoreactivity and APP levels in the astrocytes ^36,40,54^ using primary human cortical astrocytes as the experimental model. Cultures were treated with 2 μg/ml LPS for 48 hours and GFAP-positive astrocytic profiles analysed by the end of this period with the Sholl analysis, determining the number of cell processes, the sum of intersections of processes with concentric circles, the ramification indices, and the ending cell radii (Figure 1a and b, Figure S1). Treatment with LPS increased morphological complexity of astrocytes as evidenced by a significantly increased average number of processes and the sum of intersections in comparison to untreated control (Figure 1C). In addition, LPS treated astrocytes had significantly increased cellular levels of full-length APP whereas there was no change in the ratio of the average mature (processed) versus immature (newly synthetized) APP levels compared with untreated astrocytes (Figure 1d).

**Figure 1:**
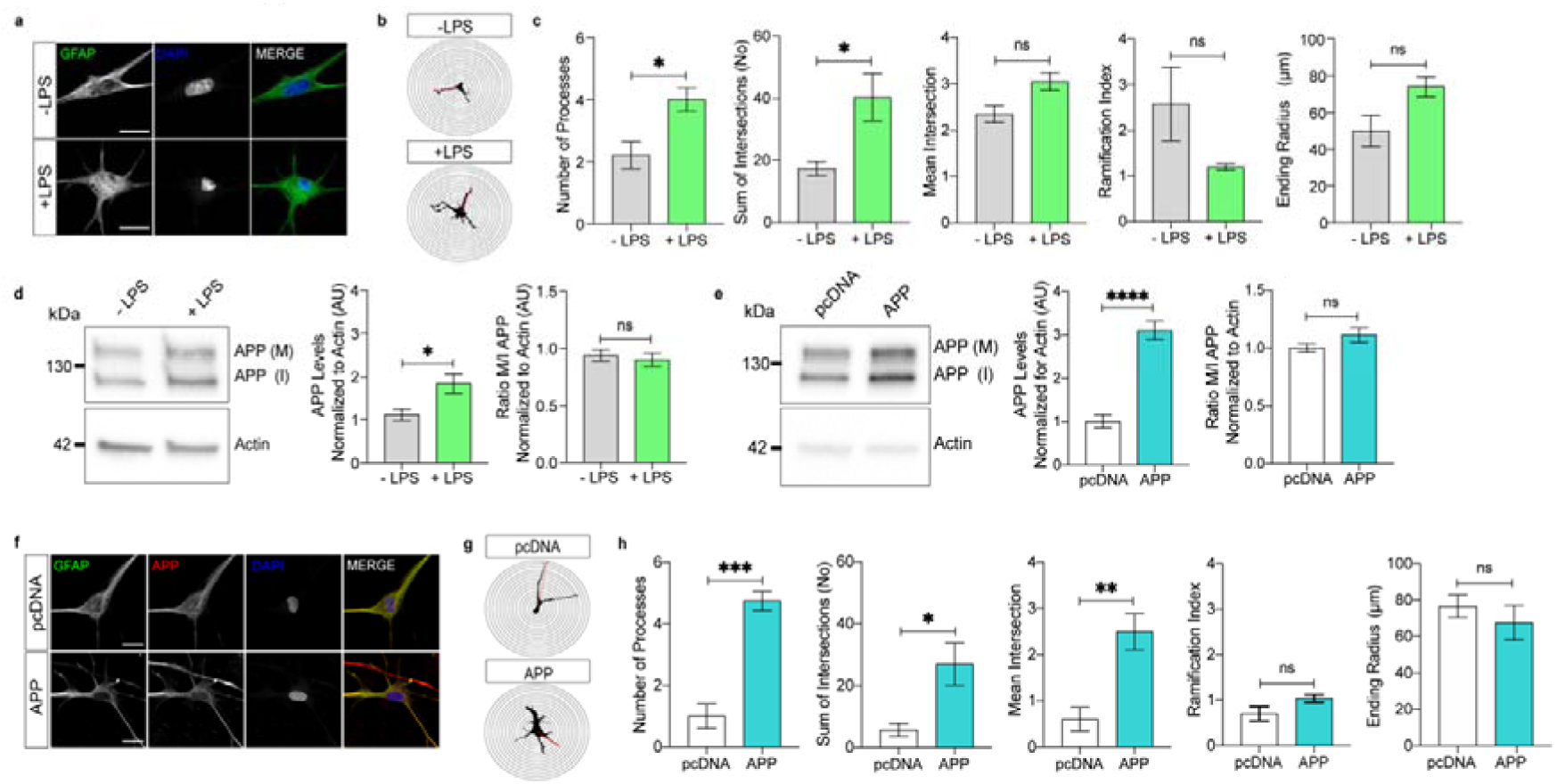
APP promotes remodelling of GFAP-positive astrocytic profiles. a. Representative images of untreated (-LPS) and LPS-treated (+LPS) primary human cortical astrocytes stained with GFAP and DAPI (Scale bar 20 μm). b. Representative image reconstructions of -LPS and +LPS GFAP-stained astrocytes based on the Sholl analysis using 5 μm spacing between the concentric circles. c. Comparison of key Sholl analysis parameters between -LPS and +LPS GFAP-stained astrocytes (5 to 8 cells per biological sample, N=3 biological samples per condition, Student t-test, * = p<0.05) d. Representative blot showing APP levels in -LPS and +LPS astrocytes normalised for actin levels as a loading control. Graphs show mean levels of APP normalised for actin (left) and the mean ratio between mature (M) versus immature (I) APP (right) (N=5 biological samples per condition, Student t-test, * = p<0.05). e. Representative blot showing APP levels in astrocytes transfected with either pcDNA3.1 (pcDNA) or APP770-pcDNA3.1 (APP) normalized for actin levels as a loading control. Graphs show average mean levels of APP normalised for actin (left) and the mean ratio of mature (M) versus immature (I) APP (right) (N=5 biological samples per condition, Student t-test, ns = not significant). f. Representative images of astrocytes transfected with either pcDNA3.1 or APP770-pcDNA3.1 and stained with GFAP, APP and DAPI (Scale bar 20 μm). g. Representative image reconstructions of GFAP-stained astrocytes transfected with either pcDNA3.1 or APP770-pcDNA3.1 based on the Sholl analysis using 5 μm spacing between the concentric circles. h. Comparison of key Sholl morphometric analysis of GFAP-stained astrocytes transfected either with pcDNA3.1 or with APP770-pcDNA3.1 (3-5 cells per biological sample, N = 4 biological samples per condition, Student t-test, * = p<0.05, ** = p < 0.01, *** = p < 0.001).

As the next step, we investigated whether overexpression of the 770 amino acid long APP (APP770-pcDNA3.1) alone induces astrocyte remodelling in the absence of stressors. Transfection of astrocytes with APP770-pcDNA3.1 resulted in a roughly 3-fold increase in the average levels of cellular APP, with no impact on the mature versus immature APP ratio compared to astrocytes transfected with the pcDNA3.1 vector (Figure 1e, Figure S2). As with LPS induction, Sholl analysis revealed that sole overexpression of APP was accompanied by a significant increase in the number of processes, sum of intersections and mean intersections of GFAP-positive astrocytes (Figure 1f-h).

Hence a canonical instigator of reactive astrogliosis (LPS) and overexpression of APP both result in the remodelling of the intermediate filament network. This signifies that an increase of intracellular APP on its own is sufficient to induce morphological features characteristic of reactive astrocytes. However, the characteristics of astrocytes are very diverse and complex and morphological changes alone are insufficient to conclude on their acquisition of a pathologically relevant reactive state. Thus, we examined next whether or not these morphological changes are accompanied by typical functional changes characteristic for reactive astrocytes.

### APP induces generation of cytokines and activates interferon pathways

To validate an experimental paradigm to track more functional changes of the astrocytes, we exposed primary human cortical astrocytes for 48 hours to LPS and then measured cytokines using a well-established ELISArray (Figure 2a). In accordance with previous reports ^32,33,35^, LPS increased cytokine levels in astrocytes. In particular, LPS led to significant increases in the levels of IFNγ, interleukin-6 (IL-6), TNFα, granulocyte colony-stimulating factor (G-CSF), and intereukin-12 (IL-12) and a significant decrease in TGFβ levels compared with unstimulated astrocytes.

**Figure 2:**
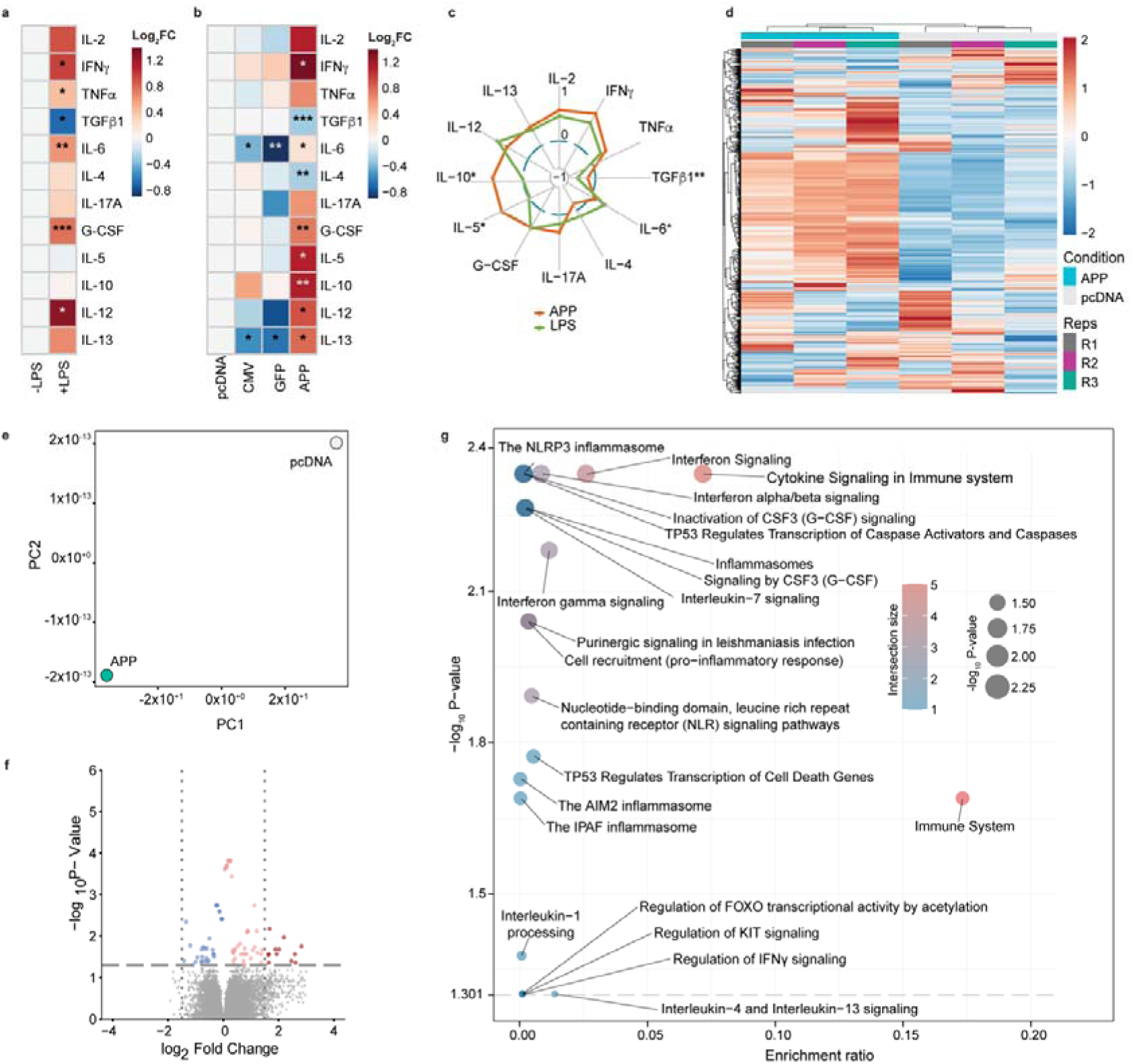
APP increases expression of cytokines in astrocytes. a. Heatmap showing changes in abundances of cytokines in -LPS and +LPS astrocytes (N = 3 biological samples per condition, Student t-test, * = p < 0.05, ** = p < 0.01, *** = p < 0.001). Individual protein concentrations (pg/mg) were log2 transformed. b. Heatmap showing changes in abundances of cytokines in pcDNA3.1 (pcDNA), CMV, GFP-pdDNA3.1 (GFP) and APP770-pcDNA3.1 (APP) transfected astrocytes (N = 3 biological samples per condition, Student t-test, * = p < 0.05, ** = p < 0.01, *** = p < 0.001). Individual protein concentrations (pg/mg) were log2 transformed. c. Radar plot showing changes in cytokine profiles between LPS treated and APP770-pcDNA3.1 transfected astrocytes (N = 3 biological samples per condition, ANOVA with Tukey post-hoc test, * = p < 0.05, ** = p < 0.01). d. Bidirectionally clustered heatmap showing the expression of 1000 genes with the highest variability. Colour annotations differentiate individual astrocyte treatments and replicates. e. Principal Component Analysis (PCA) of transcriptomic profiles of APP770-pcDNA3.1 *versus* pcDNA3.1 empty vector transfected astrocytes. f. Volcano plot showing differentially expressed (DE) genes in astrocytes transfected with APP770-pcDNA3.1 and pcDNA3.1. The horizontal line indicates statistically significantly DE genes, vertical dotted lines indicate the log 2 FC of ±1.5. g. The most significantly changed REACTOME pathways by the DE genes with log 2 FC exceeding the cut off of ±1.5.

We next measured cytokine levels in primary human cortical astrocytes transfected with pcDNA3.1 (empty vector), CMV and GFP-pcDNA3.1 as controls or with APP770-pcDNA3.1 (Figure 2b). Transfection of astrocytes with APP770-pcDNA3.1 resulted in a significant increase in IFNγ, G-CSF, interleukin-5 (IL-5), interleukin-10 (IL-10), IL-12, interleukin 13 (IL-13) and a significant decrease in IL-6, interleukin-4 (IL-4) and TGFβ1 levels compared with astrocytes transfected with pcDNA3.1. Transfection of astrocytes with CMV empty vector or GFP-pcDNA3.1 produced significant changes in IL-6 and IL-13 levels compared with pcDNA3.1 only. This indicates a marginal effect of empty vector or generally foreign protein on the cytokine expression and DNA sensing pathways ^55^. In contrast to LPS treatment, overexpression of APP produced significant changes in astrocytic levels of TGFβ1, IL-5 and IL-10 and IL-6 (Figure 2c). The results of these experiments indicate that APP overexpression induces cytokine production in astrocytes with cytokine changes mediated by APP exhibiting a unique cytokine profile that differs from the one obtained following LPS treatment.

To test whether the observed changes in cellular cytokine levels following APP overexpression are reflected in enhanced gene expression and to identify the most significantly activated cellular programs, transfected astrocytes with either pcDNA3.1 empty vector or APP770-pcDNA3.1 were isolated and sequenced for their RNA. Heatmap and principal component analysis showed significant differences in the transcriptional profiles in the APP770-pcDNA3.1 compared with the pcDNA3.1 transfected astrocytes (Figure 2d and e). Volcano plots identified several significantly upregulated and no downregulated differentially expressed genes in the APP770-pcDNA3.1 compared with the pcDNA3.1 transfected astrocytes (Figure 2f). Gene ontology enrichment analysis showed that overexpression of APP mostly upregulated interferon, cytokine and inflammasome signalling (Figure 2g). Indeed, overexpression of APP led to significantly increased astrocyte IFNγ and IFNβ at the transcriptional and protein level (Figure 2g and Figure S3). Comparison of the transcriptomic profiles between astrocytes transfected with either APP770-pcDNA3.1 or APP770-pcDNA3.1 carrying the familial Swedish, Florida, Austrian, Arctic or Icelandic APP mutation^56–60^ showed no differences between profiles, indicating that activation of interferon, cytokine, and inflammasome signalling pathways depends on the levels of full-length APP rather than on the mutation-induced changes in its proteolytic processing (Figure S4). These results indicate that overexpression of APP induces not only morphological, but also functional features of reactive astrocytes.

### Increased APP- and IFN**γ** in reactive astrocytes following TBI

To test for the physiological relevance of the observed relationship between APP and the cytokines in primary human cortical astrocytes, we studied APP- and IFNγ in reactive astrocytes following controlled cortical impact (CCI), a well-established mouse model of traumatic brain injury (TBI) (Figure 3a) ^61^. We first examined the morphology of GFAP-stained astrocytes in the corpus callosum in an area positioned just under the CCI region using the Sholl analysis (Figure 3b and c). Two months following CCI, GFAP-immunoreactive astrocytes exhibited significantly increased numbers of processes, and more intersections and mean intersections in TBI compared with control mice (Figure 3D). As previously reported,^62^ TBI triggered reactive astrogliosis, which led us next to investigate the relationship between APP- and IFNγ-immunoreactivities in reactive astrocytes following TBI. After a series of pilot experiments to optimize labelling and imaging (Figure S5), GFAP-stained reactive astrocytes showed significantly increased APP- and IFNγ-immunoreactivities in TBI compared with sham operated mice (Figure 3e and f). In addition, a significant positive correlation was observed between the mean intensities of APP-and IFNγ-immunoreactive signals in astrocytes in control and TBI mice (Figure 3g). These experiments provide *in vivo* corroboration of the interaction between APP and IFNγ previously found in cell culture by identifying a direct relationship between APP- and IFNγ-immunoreactivities in reactive astrocytes following TBI.

**Figure 3:**
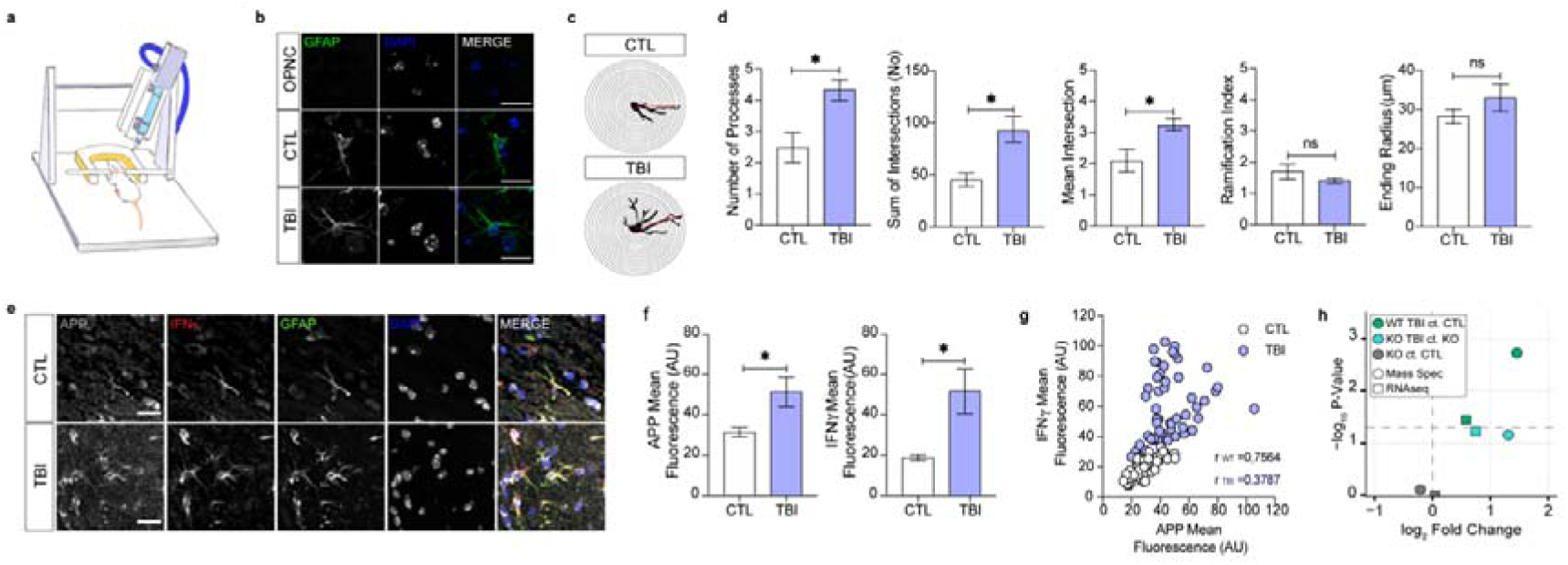
APP and IFNγ in astrocytes following TBI. a. Cartoon depicting the controlled cortical impact (CCI) injury model of the traumatic brain injury (TBI). b. Representative confocal microscopy images of OPNC (omitted primary negative control) and GFAP-stained astrocytes in the corpus callosum of control and TBI mice (scale bar 20Cµm). c. Representative reconstructed images of GFAP-immunoreactive astrocytes of the corpus callosum of control and TBI mice based on the Sholl analysis with a 5 μm interval of the concentric circles. d. Measurements of key Sholl parameters in GFAP-stained astrocytes in control and TBI mice (5 to 7 per animal, N = 4 mice, Student t-test, * = p < 0.05). e. Representative confocal microscopy images of APP- and IFNγ-immunoreactivities in GFAP-stained astrocytes in corpus callosum of control and TBI mice (Scale bar 20Cµm). f. Graph showing mean APP- and IFNγ-immunoreactivities in GFAP-stained astrocytes in the corpus callosum of control and TBI mice (N = 4 mice, Student t-test, * = p < 0.05). g. Relationship between mean APP- and IFNγ-immunoreactivities in the GFAP-stained astrocytes in the corpus callosum of control and TBI mice (N = 4 mice, Pearson correlation coeficient ANOVA two-tailed **** = p < 0.0001 and ** = p < 0.01).

Finally, we examined whether APP is essential for the *in vivo* increase in GFAP-content in astrocytes after TBI. To this end, we analysed GFAP expression and levels in wildtype mice (WT), WT mice following CCI (WT TBI), APP knock out mice (KO) and KO mice following TBI (KO TBI) obtained from an ongoing RNA-Seq and tandem mass tag mass spectrometry (TMT-MS) study. Compared with brains from WT mice, GFAP mRNA expression and protein levels in WT TBI mice were significantly (N=3; 0.58-fold (*p*=0.035) and 1.45-fold (*p*=0.002)) higher as revealed by RNA-Seq and TMT-MS, respectively (Figure 3h). In contrast, there were no significant differences in GFAP levels, determined by RNA-Seq or TMT-MS when comparing KO TBI with KO (N=3; *p*-RNA-Seq = 0.059, TMT-MS = 0.069) or KO to WT (N=3; *p*-RNA-Seq = 0.938, TMT-MS = 0.744). These findings align with our previous observations in cell culture, indicating an *in vivo* connection between APP- and IFNγ-immunoreactivities in reactive astrocytes following TBI and underscore the crucial role of APP in triggering reactive astrogliosis.

### LPS-induced reactive astrogliosis requires APP

Our findings suggest that APP induces morphological and functional features characteristic of reactive astrocytes in cell culture as well as in the *in vivo* brain. These experiments, however, do not critically test whether APP is required for reactive remodelling of astrocytes. To test this hypothesis, we deleted APP and treated cultured astrocytes with LPS and subsequently analysed morphology of GFAP-positive astrocytic profiles and measured levels of a panel of IFN-stimulated gene (ISG) products. To confirm the impact of LPS treatment and the deletion of APP induced by the multiMIR-APP lentiviral vector, we initially measured cellular APP levels normalized to actin using western blots (Figure 4a). This analysis revealed a significant increase and decrease in APP levels in astrocytes subjected to LPS treatment or transduced with the multiMIR-APP lentiviral vector, respectively. Whether we used a multiMIR-APP vector or a control vector (multiMIR-Scramble), Sholl analysis of GFAP-labelled primary human cortical astrocytes (Figure 4b-c) demonstrated similar cell morphologies. Consistent with our previous findings, LPS-treated astrocytes displayed a notable increase in the mean number of processes compared to untreated astrocytes (Figure 4d). Deletion of APP using the multiMIR-APP lentiviral vector nullified the LPS-induced increase in the mean number of processes in astrocytes. Consequently, astrocytes lacking APP were unable to mount a reactive response to LPS.

**Figure 4:**
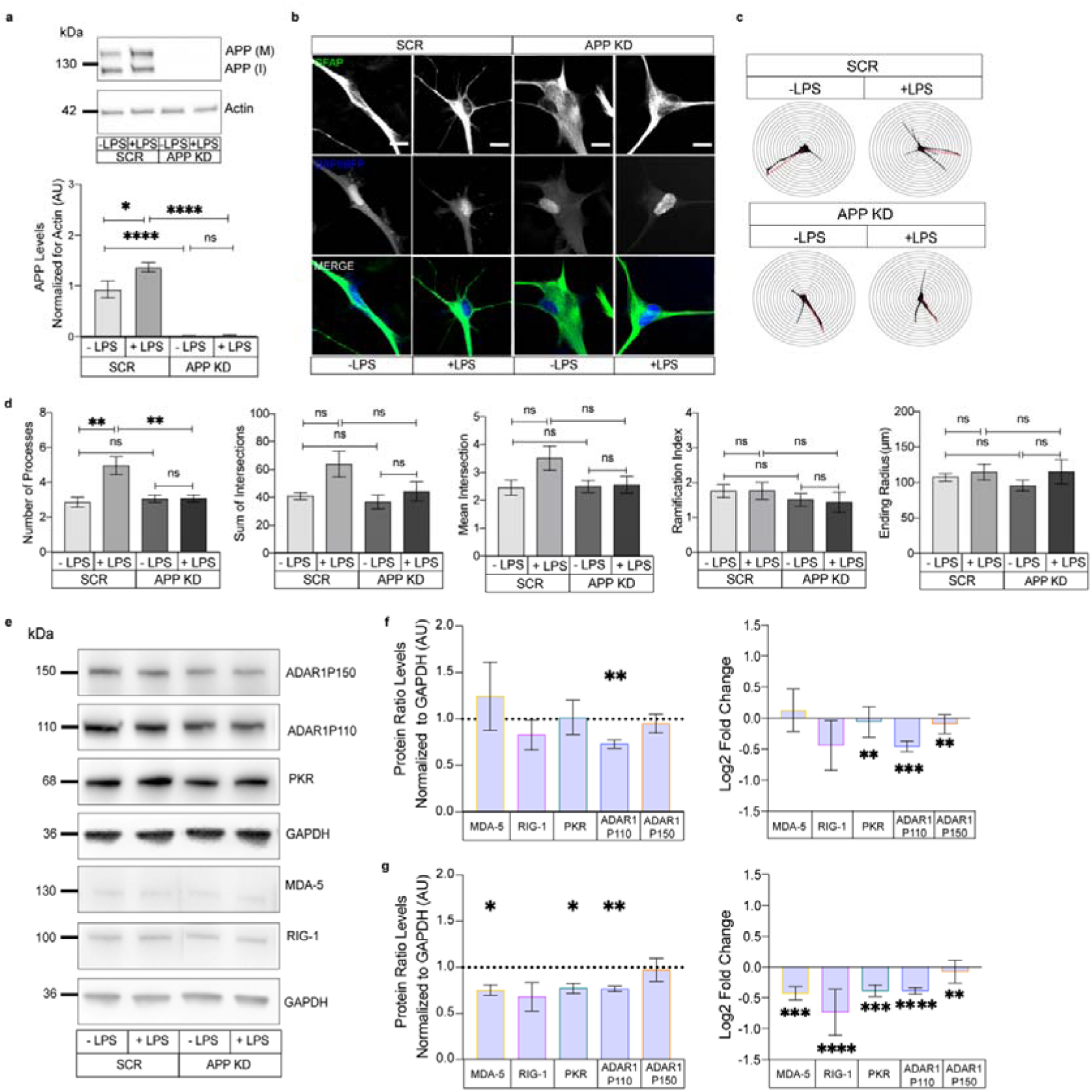
Deletion of APP hampers reactive astrogliosis in response to LPS. a Representative western blot showing levels of APP normalised for actin as a loading control in untreated (-LPS) and LPS treated (+LPS) astrocytes transduced with multiMIR-Scramble (SCR) and multiMIR-APP (APP KD) lentiviral vectors. Graph showing mean APP levels normalized for actin (N = 5 biological replicates, ANOVA with Tukey’s multiple comparisons test, ns = not significant, * = p < 0.01, **** = p < 0.01). b. Representative confocal microscopy images of GFAP-stained -LPS and +LPS astrocytes transduced with multiMIR-Scramble and multiMIR-APP lentiviral vectors (Scale bar 20Cµm). c. Representative reconstructed images of GFAP-stained -LPS and +LPS astrocytes transduced with multiMIR-Scramble and multiMIR-APP lentiviral vectors based on the Sholl analysis with a 5 μm interval of the concentric circles. d. Measurements of key Sholl parameters in GFAP-immunoreactive -LPS and +LPS astrocytes transduced with multiMIR-Scramble and multiMIR-APP lentiviral vectors (10 to 27 cells per treatment, N = 4 biological replicates/mice, ANOVA with Tukey’s multiple comparisons test, ** = p < 0.01). e. Representative western blot of ISG product levels and GAPDH levels as a loading control in -LPS and +LPS astrocytes transduced with multiMIR-Scramble and multiMIR-APP lentiviral vectors. f. Graph showing ratios of GAPDH normalised cellular ISG product levels (left, One Sample t-test, * = p < 0.05, ** = p < 0.01) and their geometrical differences presented as log2 fold changes (right, Student t-test with Benjamin-Hochberg correction, ** = p < 0.01, *** = p < 0.001) between multiMIR-Scramble and multiMIR-APP transduced astrocytes (N = 4 biological replicates). g. Graph showing ratios of GAPDH normalised cellular ISG product levels (left, One Sample t-test, * = p < 0.05, ** = p < 0.01) and their geometrical differences presented as log2 fold changes (right, Student t-test with Benjamin-Hochberg correction, *= p < 0.01, **= p < 0.01, *** = p < 0.001 **** = p < 0.0001 between -LPS and +LPS multiMIR-Scramble and multiMIR-APP transduced astrocytes (N = 4 biological replicates).

We next measured cellular levels of a panel of ISG products including adenosine deaminase acting on RNA p110 (ADAR1p110), adenosine deaminase acting on RNA p150 (ADAR1p150), protein kinase R (PKR), melanoma differentiation-associated protein 5 (MDA-5) and retinoic acid-inducible gene 1 (RIG-1) all normalised to GAPDH (Figure 4e). To examine changes in the levels of cellular ISG products following APP deletion and LPS-treatment, we measured either their expression ratios or log_2_ fold changes, first between astrocytes transduced with multiMIR-APP versus multiMIR-Scramble, and then between LPS-treated astrocytes transduced with multiMIR-APP versus multiMIR-scramble lentiviral vectors. We observed significant reduction in ADAR1p110- and PKR-ratio and significant log_2_ fold change of both ADAR1 isoforms between multiMIR-APP and multiMIR-Scramble transduced astrocytes (Figure 4f). Comparing cellular ISG product levels between LPS-treated multiMIR-APP- and multiMIR-Scramble-transduced astrocytes, we observed significant reductions in MDA5, PKR and ADAR1p110 ratios and in MDA5, RIG-1, PKR and both ADAR1 isoforms log2 fold changes (Figure 4g). The results of these measurements indicated that APP deletion alone perturbs levels of ISG products in naïve astrocytes and prevents LPS-induced increase in ISG products. Collectively, these data indicate that APP deletion suppresses reactive astrogliosis.

## DISCUSSION

Increased APP levels in reactive astrocytes were widely reported ^32,36,37,39,40,63^, but the underlying rationale for increased APP levels in reactive astrocytes remained so far largely veiled remained so far largely veiled. We here demonstrate that increased expression of APP translates into remodelling of GFAP-positive astrocytic morphology profiles reflecting changes in intermediate filament network, increased production of cytokines and activation of the IFN pathways. An increase in APP alone therefore is sufficient to induce reactive astrogliosis. Conversely, depletion of APP abrogates LPS-induced reactive astrogliosis and reduces expression of ISG products. Our results support the idea that APP is a key player in triggering and regulation of reactive astrogliosis because suppressing APP expression is sufficient to prevent astrocytes from acquiring reactive response to stressors. Thus, APP acts as a molecular inducer and regulator of reactive astrogliosis.

To examine whether our findings apply to realistic pathological contexts, we build upon previous work in animal models of brain trauma ^22,64–67^. These previous studies, however, did not show the interaction between APP and IFN in reactive astrocytes revealed by the present study. To translate our cell culture findings into pathophysiologically relevant animal models, we confirmed an association between APP and IFN levels in reactive astrocytes affected by TBI trauma ^61^. Understanding this relationship as well as the crosstalk between APP and cytokines requires further validation and mechanistic studies, perhaps by taking advantage of human stem cell derived astrocyte cell lines ^68–70^ in addition to the transcriptomic and proteomic analyses of individual astrocytes derived from astrocyte-centred animal models and human post-mortem brains ^71,72^.

Lack of changes in the transcriptomic profiles of astrocytes transfected with wildtype or familial APP mutations corroborates previous reports indicating that proteolytic processing of APP differs significantly between astrocytes and neurons ^73–77^ and suggests that full-length APP rather than its proteolytic fragments instigate reactive astrogliosis. Whether this is related to the dominance of the Ox-2 immunoglobulin- and Kunitz-type protease inhibitor domains-containing 770 amino acid long APP isoform ^31,38,78–80^, which is enriched in astrocytes in AD ^81,82^, remains to be investigated. A wealth of data documents that exposure to Aβ alone induces reactive astrogliosis ^83,84^ and activates IFN and other pro-inflammatory pathways ^85^. Since at least some of the morphological and functional features characteristic of reactive astrocytes can be triggered by Aβ ^86–88^, these data indicate that APP and its proteolytic fragments, such as Aβ, may regulate the structure and function of the astrocytes in an independent as well as in complementary ways. Since APP and its Aβ fragment have been demonstrated to induce IFN signalling pathways ^85^, which also regulate expression and proteolytic processing of APP including generation of Aβ ^89,90^, these studies suggest a putative positive feedback or a feed-forward loop between APP, its proteolytic fragments and the IFN pathways ^54,91,92^. It is plausible to hypothesize that such a positive feedback loop plays a role in IFN-induced remodelling of the intermediate filament network in the reactive astrocytes^93–98^. Emerging data showing that APP binds to and modifies regulators of cytokine expression ^99–102^, together with reportedly blunted innate immune response observed in mice deficient in APP^103^, also suggest that consequences of the interplay between APP and the immune networks might be significantly more far-reaching than previously thought ^104^.

In conclusion, our experiments reveal a role of APP as an instigator of reactive astrogliosis. By manipulating APP expression, we observed changes in astrocyte morphology and function, implicating APP as a molecular regulator of reactive astrogliosis. Our study enhances the understanding of the intricate relationship between APP and astrocytic reactivity with extensive implications in furthering our understanding of brain pathologies and developing novel therapies against neuroinflammatory diseases. Given the accumulating evidence that reactive astrocytes are neuroprotective ^15,105,106^, our experiments further strengthen the previously proposed neuroprotective role of APP across different CNS cell types. Without doubt, astroglial APP plays a key role in the neural tissue response to damage. In prospect, our findings merit to be explored in genetic, demyelinating and other brain disorders which involve reactive astrogliosis ^108–110^.

## MATERIALS AND METHODS

### Cell cultures

Human primary cortical astrocytes (HA, ScienCell) were grown according to manufacturer’s instructions with two modifications. Astrocyte medium (#1901, ScienCell) was not supplemented with fetal bovine serum (FBS) and the poly-L-lysine was replaced with the matrigel matrix (#356231, Corning). HA were cultured at 37°C and 5% CO_2_. To ensure reproducibility, all experiments were performed with HA at passage number 5. Whenever experiments required treatment with a stressor, the astrocytes were incubated with 2μg/ml of bacterial endotoxin LPS (LPS-EB, Invitrogen) for 48 hours.

### Transfection

HA at 70% confluency were transiently transfected using Lipofectamine 2000 following manufacturer’s instructions (#12566-014, Thermo Fisher). The construct consisting of human APP770 cDNA in the pcDNA3.1 backbone was custom made (GenScript, Piscataway) and then sequenced. The plasmid was transfected into HA in a 1:2 DNA to Lipofectamine 2000 ratio. HA were lysed or fixed 48 hours later.

### Transduction

To knock-down APP, HA were transduced with a lentivector expressing a multimiRNA system linked to a blue fluorescent reporter protein (Flashtherapeutics, Toulouse), which carried either a multiplex of 6 miRNAs targeting APP (multiMIR-APP, pV.2.3.2029) or scramble sequences of the siRNAs targeting APP sequences (multiMIR-Scramble, pV.2.3.2028). The multiMIR-Scramble and multiMIR-APP lentiviral vectors were transduced into astrocytes using 5, 10 or 20 multiplicity of infection (MOI). 5 days after transduction, the cells were lysed, and APP levels measured by western blot. MOI 10 was selected for the experiments as it showed no APP signal compared with APP levels measured in astrocytes transduced with the multiMIR-Scramble lentiviral vector.

### Animal models

3-month-old male wildtype mice (C57Bl/6, Jackson Laboratories, Bar Harbor) were subject either to sham surgery or controlled cortical impact (CCI) using a well-established protocol^61^. All mice were euthanized by rapid decapitation and brains collected for the analyses. All animal use protocols were approved by the Veterans Administration San Diego Healthcare System Institutional Animal Care and Use Committee (San Diego, USA). Mice were handled in compliance with the Guide for the Care and Use of Laboratory Animals (National Academy of Science, Washington, D.C).

### Tissue sampling

Sham operated and CCI exposed wildtype mouse brains were fixed in 4% PFA, embedded in tissue freezing medium (OCT, #14020108926, Leica Biosystems) and snap-frozen using cooled isopentane. Frozen samples were stored at -80°C until sectioning. Fixed brains were later cut horizontally at 10-micron thickness using a Leica CM1950 cryotome.

### Western blotting

Cells were washed with cold phosphate-buffered saline (PBS), scraped off the plates and lysed on ice for 30 min with 1X RIPA buffer (#20-188, Merck) supplemented with phosphatase (#P5726, Merck) and proteinase inhibitors (#P8340, Merck). The lysates were then span down (20 min, at 4 °C, 14.000 RPM) and protein concentration of collected supernatants measured using the Pierce BCA Protein Assay Kit (#323227, Thermo Scientific, 2322<u>7</u>). The samples adjusted to contain equal amounts of protein (10 μg or 12 μg) were next denatured (5 min at 95°C) in loading buffer (#1610747, BioRad) and run on 10% TGX (#4561034, BioRad) or mPAGE 4-12% Bis-Tris Precast gels (#MP41G10, Merck). Separated proteins were then transferred to PVDF or Nitrocellulose membranes according to manufacturer’s instructions (Bio-Rad, Hercules). After 1 hour of blocking in 5% (w/v) BSA (bovine serum albumin) dissolved in TBS-T (20 mmol/L Tris, pH 7.4, 65 mmol/L NaCl, 0.1% Tween 20), the membranes were probed with primary antibodies against APP (1:1000, #ab3261, Abcam), β-actin diluted (1:2000, #A2228, Sigma), ADAR1 diluted (1:3000, #ABIN2855100, Antibodies-online), RIG-1 diluted (1:1000, #3743, Cell signalling), MDA5 diluted (1:800, #5321, Cell signalling), PKR diluted (1:2000, #ab184257, Abcam) and GAPDH diluted (1:50000, #60004-1-lg, Proteintech), washed in TBS-T and incubated with horseradish peroxidase–conjugated anti-rabbit (1:2000, #7074P2, Cell Signalling or 1:80000, #A0545, Merck) and anti-mouse (1:2000, #7076P2, Cell Signalling or 1:5000, #331430, Thermo Fisher) secondary antibodies. The blots were developed using ECL (BioRad, Clarity, 1705060) and captured bands analyzed using Chemidoc (BioRad, Hercules) and ImageJ ^111^

## ELISA

Cell lysates were homogenized with 1X RIPA on ice for 30 minutes and then centrifuged for 20 minutes at 14,000 g at 4 C. Cytokines were measured using a multi-analyte ELISAarray according to the manufacturer’s instructions (#MEH-003A, Qiagen). The INF-β and INF-γ were measured using single target ELISAs (#DIF50C and #DY814-05, R&D Systems).

### Immunocytochemistry

HA were cultured on coverslips, washed with PBS, fixed with 4% PFA in PBS for 30 min at RT, washed with PBS and then incubated with the blocking solution (10% goat serum, 0.1% Triton X-100 in PBS) for 1 hour at RT. Cultures were then stained overnight at 4 °C with primary antibodies against GFAP (1:500, #Z0334, Dako) and APP (1:500, #MAB348, Sigma) dissolved in antibody solution (0.1% Triton X-100 in PBS). Cells were thereafter washed with PBS and stained with secondary antibodies (#A-21206 and #A-31570, Life Technologies) for one hour at RT. Cells were then washed, stained with DAPI and mounted on slides with Mowiol. Stained cells were examined with a LSM confocal microscope (LSM780, Zeiss) using a 40x oil-immersion objective. Images were acquired and processed using the ImageJ software ^111^

### Immunohistochemistry

Mouse brain sections were blocked with MOM solution (#MKB-2213, Vector Labs,) and then incubated overnight with primary antibodies against INFγ (1:100, #MCA1301, Bio-Rad), GFAP (1:1000, #ab53554, Abcam) and APP (1:200, #ab2072, Abcam) or against Aβ (MOAB-2 1:1000, #ab126649, Abcam). The next day, the sections were incubated with donkey anti-mouse (AF555, 1:500, #A3157, Thermo Fisher), donkey anti-goat (AF488, 1:500, #A11055, Thermo Fisher) and donkey anti-rabbit (AF647, 1:500, #A31573, Thermo Fisher) or donkey anti-mouse (AF555, 1:500, #ab150106, Abcam) secondary antibodies followed by DAPI staining and Mowiol mounting.

Sections stained with secondary antibodies only were used as negative controls. Additionally, bleed through control stainings were incubated with all three primary antibodies and one secondary separately. Stained mouse brain sections were imaged using a 10x objective on AxioScan.Z1 slide scanner microscope (Zeiss, Oberkochen). To better visualize the distribution of specific markers the slides were imaged also with a 63x oil-immersion objective on a LSM780 confocal microscope (Zeiss, Oberkochen).

### Protein Extraction and quantification

Brain tissue was weighed and homogenized in lysis buffer (RIPA 1X Millipore, 20-188, 10 µl of buffer per 10 mg of tissue) supplemented with protein and protease inhibitors without detergents followed by incubation on ice for 45 minutes and centrifugation at +4°C for 20 minutes. Supernatants were collected and stored at -80°C until used for experiments. Protein concentration was determined via BCA assay (Pierce™ BCA, 23225).

### TMT-based Quantitative Mass Spectrometry

Quantitative mass spectrometry (MS) analysis was performed on brain samples from 3 animals per group by the Proteomics Core Facility EMBL, Heidelberg, Germany. MS has been carried out independently for a specific project and consequently the data will be published independently. Hence, solely GFAP data was selectively extracted for this particular objective.

Protein samples underwent isobaric labelling according the manufacturer’s instructions (TMT6plex Isobaric Label Reagent, ThermoFisher) before quantitative LC-MS/MS via UltiMate 3000 RSLC nano LC system (Dionex) as previously described (Dayon et al. 2008). The outlet of the analytical column was coupled directly to an Orbitrap Fusion™ Lumos™ Tribrid™ Mass Spectrometer (Thermo) using the Nanospray Flex™ ion source in positive ion mode. Full mass scan (MS1) was acquired with mass range 375-1500 m/z in profile mode in the orbitrap with resolution of 60000. Data dependent acquisition (DDA) was performed with the resolution of the Orbitrap set to 15000. The raw output files of IsobarQuant (protein.txt – files) were processed using the R programming language. Only proteins that were quantified with at least two unique peptides in at least 2 out of three replicates were considered for the analysis. Contra and Ipsi samples were treated separately. Proteins were tested for differential expression using the Limma R package. The FDR adjusted P-values were then calculated separately using the fdrtool R package and the lower of the two P-values was selected. A protein was annotated as differentially expressed with a negative log10 FDR adjusted P-value greater than 1.301 (corresponding to an FDR adjusted P-value < 0.05).

### Brain immunochemistry control experiments

Considering we stained mouse brains with three primary antibodies, we carried out a series of control experiments to rigorously test for possible confounding effects stemming from the contemporary use of multiple primary antibodies:

#### A. Testing for channel bleed-through

To probe for possible bleed-throughs between the microscopy channels used to visualize signals from three primary antibodies, respectively, we stained brain sections with all three primary antibodies and at the same time individually with each of the secondary antibodies and then analyzed spectral image lambda stacks to establish the emission spectra of each fluorophore and assess for possible spectral overlaps. The optimal image acquisition settings for the immunochemistry were defined based on careful assessment of the emission ranges during these control experiments, which showed no significant spectral overlap between the fluorophores.

#### B. Testing for cross-reactivity of secondary antibodies

Applying the image acquisition settings defines based on the analysis of the spectral image lambda stacks, we imaged brain sections stained with all primary antibodies and at the same time individually with each of the secondary antibodies to test for possible cross-reactivity of secondary with primary antibodies. Measurements of mean fluorescence of All individual secondary antibody conditions showed significant increase in mean fluorescence only with their corresponding and no other primary antibody.

#### C. Testing cellular distributions of different fluorophore signals

The BIOP version of JACop (Just Another Colocalization Plugin) from the ImageJ was employed to analyze cellular distributions between different fluorophore signals, which could indirectly indicate presence of possible channel bleed-through or secondary antibody cross-reactivity. Calculation of Pearsońs coefficient was used to test for relationships between cellular distributions of different fluorophore signals.

### Sholl cell morphometric analysis

Cells included in the study were selected based on a systematic random selection based on the random number table. Moreover, only cells that showed clear nuclei and were fully recognisable were included into the study. Scholl analysis of astrocytes from the *in vitro* and *in vivo* studies was then performed via the Scholl plug in ImageJ as previously described (Ferreira et al., 2014). In brief, Z-stack confocal images immunostained for GFAP were serially stacked and projected maximally. The maximal projection images of the GFAP signal in gray scale were adopted for the analysis. The plugging allowed drawing serial concentric circles with 5-μm intervals from the centre of the soma to the end of the most distant process. The starting concentric radius started where centre of the cell soma ended for each sampled astrocyte. Intersections were determined as points where the astrocytic processes cross the concentric ring. The following parameters were selected for our study:

a. Processes: number of processes originating directly from the cell soma.
b. Sum of intersections: the total sum of crossings spotted on process sections.
c. Mean intersections: the average number of intersections that occur across all the sampled radii covering astrocyte branches.
d. Ramification index: ratio between the maximum number of intersections and the number of processes.
e. Enclosing radius: refers to the distance from the centre of a cell to the furthest outer region of the same cell.

### RNA Sequencing

RNA sequencing was performed at Novogene (Cambridge, UK). Prior to Illumina sequencing, the RNA underwent a thorough quality check and assessment for degradation via Agilent Bioanalyzer. The sequencing libraries were prepared with the NEBNext® UltraTM RNA Library Prep Kit (New England BioLabs, Ipswich). The libraries were sequenced using the Illumina HiSeqTM 2000 platform.

### Bioinformatics

Differential expression (DE) analysis of fragments per kilobase of transcript per million mapped reads (FPKM) normalized gene expression data was performed using the limma R package. A negative log 10 FDR adjusted P-value > 1,301 was set as the threshold for significant DE. GO enrichment analysis of the DE genes with log 2-fold change greater than ±1.5 was implemented using the gProfiler web engine. A P-value of 0.05 was selected as the threshold for deciding whether a gene set was significantly enriched.

### Statistical analysis

Data are presented as mean ± SEM. Differences between the two groups were analyzed using t-test. Multigroup differences were assessed by ANOVA with Tukey post hoc test. All statistical analyses were performed as two-tailed with the exception of Pearson’s correlation (one-tailed) and all the p< 0.05 were considered statistically significant. Data analysis and visualizations were performed in RStudio (v.2022.07.2 with R environment v.4.2.1) as well as in Graphpad Prism 9 (La Jolla, USA).

## ABBREVIATIONS

APP: Amyloid Precursor Protein
GFAP: glial fibrillary acidic protein
IFN: interferon
ISG: IFN stimulated gene
LPS: lipopolysaccharide
TBI: traumatic brain injury

## ACKNOWLEDGEMENTS

We are most thankful for the continues feedback by Drs Clara Limbäck-Stokin and Kenneth Moya. We thank current and past members of the Stokin Lab for support and feedback.

## FUNDING

The completion of this research project would not be possible without the support from the European Union: Next Generation EU – Project National Institute for Neurological Research (LX22NPO5107 (MEYS)) (G.B.S), the European Regional Development ENOCH grant CZ 02.1.01/0.0/16_019/0000868 (G.B.S), the European Regional Development MAGNET grant CZ.02.1.01/0.0./15_003/0000492 (G.B.S.), the P30 Centre Core grant AG062429 (R.R.), the Slovenian Research Agency Core Research Program P3 310 Cell Physiology (R.Z.) and the Czech Science Foundation GAČR grant 21-27329X (M.O.C.).

## DATA AVAILABILITY STATEMENT

The authors confirm that the data supporting the findings of this study are available within the article and or its supplementary materials.

## SUPPLEMENTARY FIGURES AND LEGENDS

**Figure S1:**
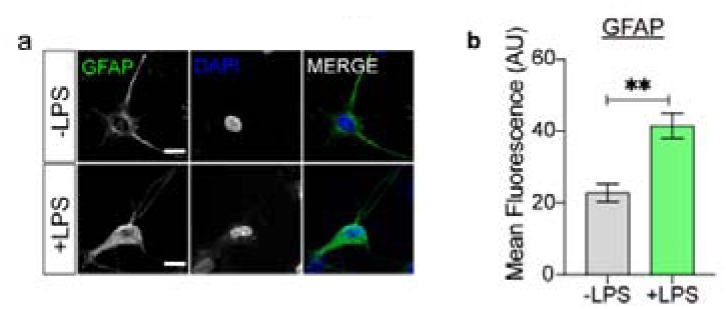
GFAP-immunoreactivity of untreated and LPS treated astrocytes. Representative images (a) and mean fluorescence (b) of -LPS and +LPS GFAP-stained primary human cortical astrocytes (N=3 biological samples per condition, Student t-test, ** = p<0.01).

**Figure S2:**
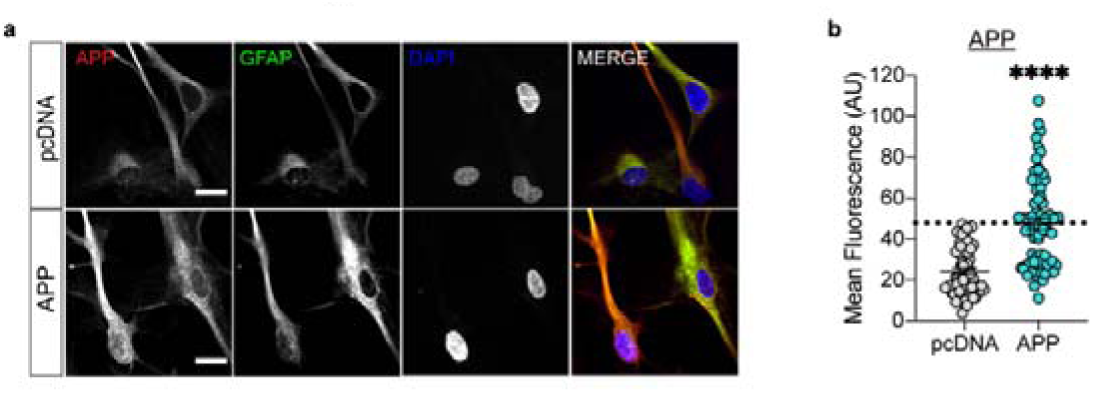
APP transfection efficiency. a. Representative images of astrocytes transfected with pcDNA3.1 (pcDNA) or APP770-pcDNA3.1 (APP) labelled with APP, GFAP and DAPI (Scale bar 20 μm). b. Average efficiencies of pcDNA3.1 or APP770-pcDNA3.1 transfection in astrocytes, dotted horizontal line shows mean fluorescence values above the ones found in pcNDA3.1 empty vector transfected astrocytes (N = 4 biological replicates, Student-test, **** = p <0.0001).

**Figure S3:**
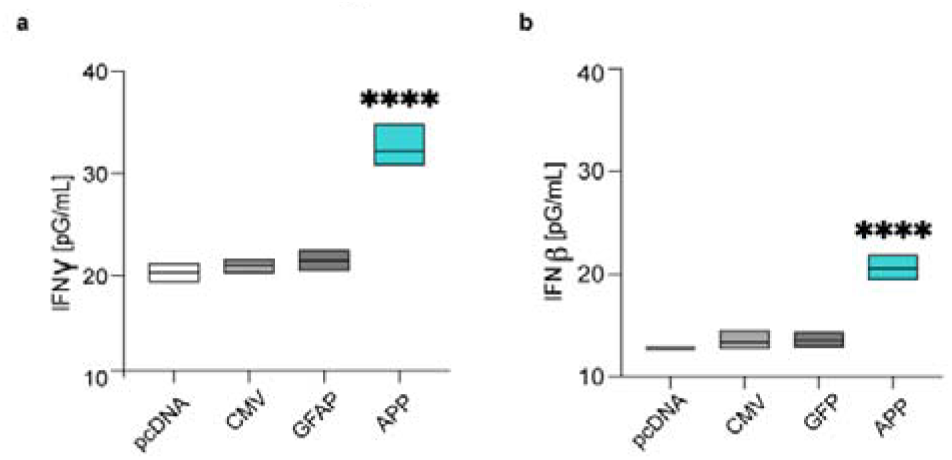
Measurements of cellular IFNγ and IFNβ levels. Cellular levels of IFNγ (A) and IFNβ (B) in astrocytes transfected with pcDNA3.1, CMV, GFP-pcDNA3.1 and APP770-pcDNA3.1 (N = 3 biological replicates per condition, ANOVA with Dunnet post-hoc test, **** = p < 0.0001).

**Figure S4:**
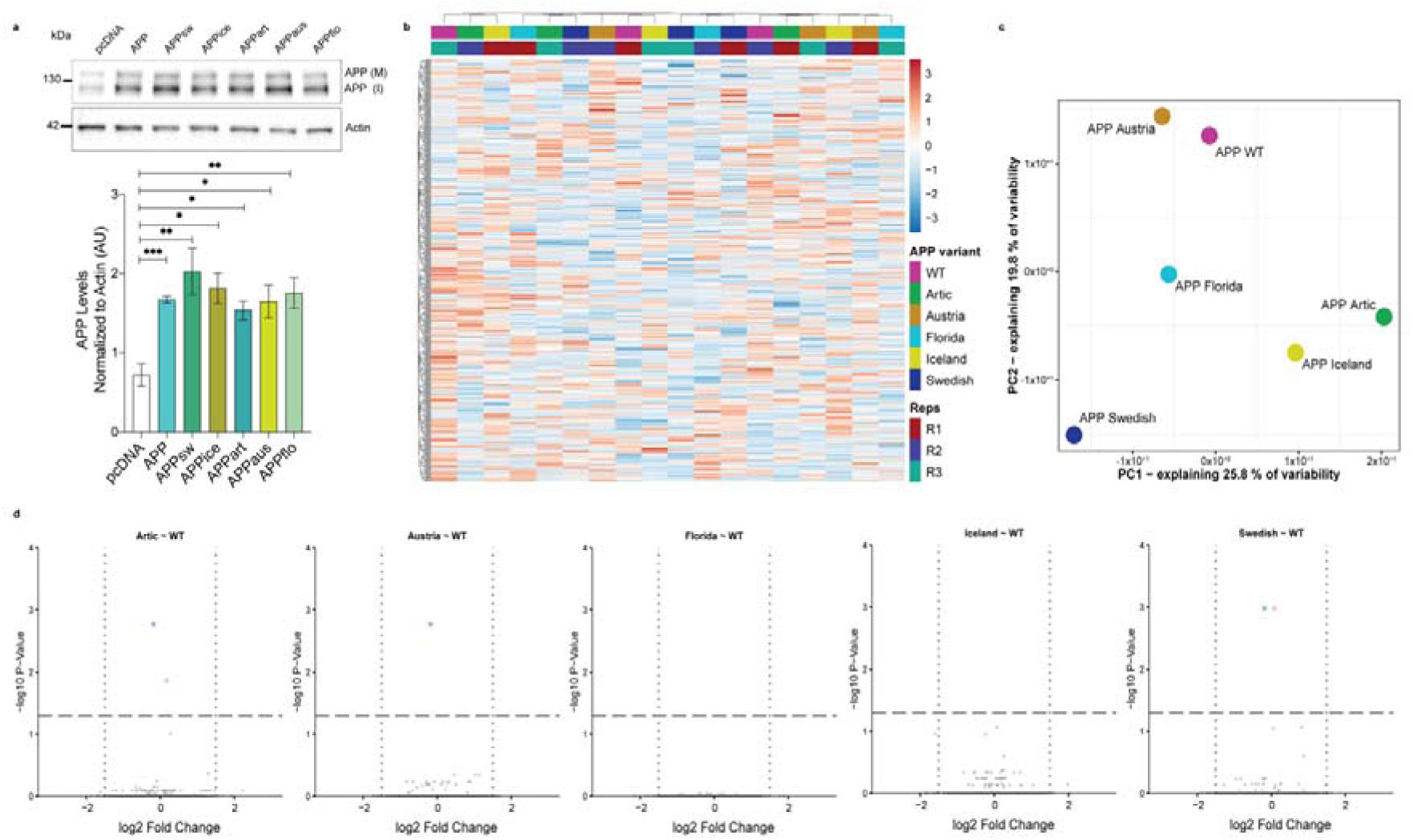
Familial APP mutations do not affect APP induced transcriptional profile of the astrocytes. a. Representative western blot of APP and actin (loading control) levels in astrocytes transfected with pcDNA3.1 (pcDNA), APP770-pcDNA3.1 (APP) or APP770-pcDNA3.1 carrying the Swedish (APPsw), Icelandic (APPice), Artic (APPart), Austrian (APPaus), Florida (APPflo) mutation correspondently. Graph showing average APP levels normalized for actin (N=3 biological replicates per condition, ANOVA with Dunnet post-hoc test, * = p < 0.05, ** = p < 0.01, *** = p < 0.001). b. Bidirectionally clustered heatmap displaying the expression of top 1000 genes with the highest variability. Colour annotations differentiate individual astrocyte treatments and individual replicates. c. Principal Component Analysis (PCA) of transcriptomic profiles of APP770-pcDNA3.1 or APP770-pcDNA3.1 carrying the different mutations shows clear differences among each group. d. Volcano plots showing DE genes in astrocytes transfected with APP770-pcDNA3.1 or APP770-pcDNA3.1 carrying one of the different mutations studied. The horizontal line indicates statistically significantly DE genes, vertical dotted lines indicate log 2 FC of ±1.5.

**Figure S5:**
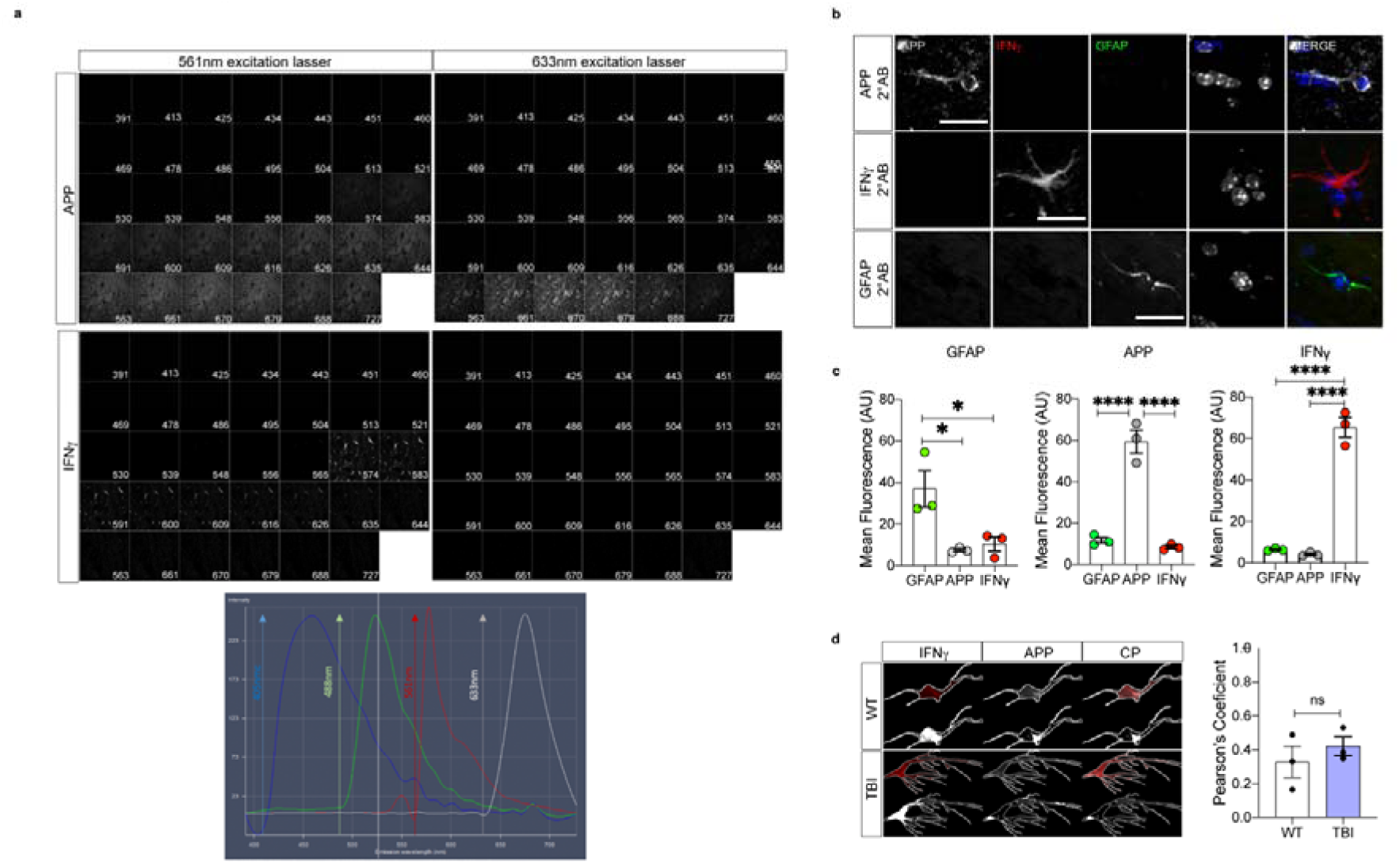
Brain immunochemistry control experiments. a. Analysis of the APP, IFNγ, GFAP and DAPI spectral image lambda stacks obtained from brain sections stained with all primary antibodies and at the same time with each of the secondary antibodies separately to establish individual fluorophore emission wavelength spectra. Following 405, 488, 561 and 633 nm excitation, there is no significant overlap between the emission wavelength spectra of different fluorophores. b. Representative images of brain sections stained with all primary antibodies and at the same time with each of the secondary antibodies separately under the optimal image acquisition settings defined based on the analysis of the spectral image lambda stacks (as shown in A, scale bar 20 μm). c. The mean fluorescence signals derived from APP, IFNγ and GFAP channels brain sections stained with all primary antibodies and at the same time with each of the secondary antibodies separately demonstrating no significant cross-reactivity of secondary with primary antibodies. (3-5 cells per mouse brain, N=3 mice, ANOVA with Dunnet post-hoc test, * = p < 0.05, **** = p < 0.0001). d. Representative images of the JACop-based co-localization analysis in GFAP-stained astrocytes showing thresholded mask of APP and IFNγ signals as well as the APP and IFNγ signal colocalization pixels (CP) in control and TBI mice (top images show APP and IFNγ signals in red and grey scale, bottom images show all pixels in white). Graph showing Pearsońs coefficients indicating weak relationship in the cellular distribution of APP and IFNγ signals and no differences between control and TBI mice (3-5 cells per mouse brain, N=3 mice, ns = not significant).

## AUTHORS CONTRIBUTIONS

Conceptualization, G.B.S.;

Methodology, G.V.J., I.G.O., D.V., C.A., K.T., S.W. K.L.K., V.L.S., M.Č., B.P.H., D.H., M.M., R.Z., L.K., M.O., R.R. and G.B.S.;

Investigation, G.V.J., I.G.O., D.V., C.A., K.T., S.W., K.L.K., V.L.S., M.Č., B.P.H., D.H., M.M. and G.B.S.;

Formal Analysis, G.V.J., I.G.O., D.V., J.S.N., K.L.K., D.H. and G.B.S.;

Data Curation, G.V.J., J.S.N. and G.B.S.;

Writing, G.V.J. and G.B.S.;

Revisions, G.V.J., D.H., R.Z., A.V., L.K., M.O., R.R. and G.B.S.;

Supervision, G.B.S.;

Project Administration, G.B.S.;

Funding Acquisition, G.B.S.;

## Notes

### Competing Interest Statement

The authors have declared no competing interest.

